# Blunted glucocorticoid responsiveness to stress causes behavioral and biological alterations that lead to posttraumatic stress disorder vulnerability

**DOI:** 10.1101/2023.08.20.554001

**Authors:** Silvia Monari, Isabelle Guillot de Suduiraut, Jocelyn Grosse, Olivia Zanoletti, Sophie E. Walker, Michel Mesquita, Tobias C. Wood, Diana Cash, Simone Astori, Carmen Sandi

**Affiliations:** Laboratory of Behavioral Genetics, Brain Mind Institute, School of Life Sciences, Ecole Polytechnique Fédérale de Lausanne, 1015 Lausanne, Switzerland; Department of Neuroimaging, Institute of Psychiatry, Psychology and Neuroscience, King’s College London, London, UK

**Keywords:** PTSD, glucocorticoids, hippocampus, fear extinction, REM sleep, norepinephrine

## Abstract

**BACKGROUND:** Understanding why only a subset of trauma-exposed individuals develop posttraumatic stress disorder (PTSD) is critical for advancing clinical strategies. A few behavioral (deficits in fear extinction) and biological (blunted glucocorticoid levels, small hippocampal size, and rapid-eye movement sleep (REMS) disturbances) traits have been identified as potential vulnerability factors. However, whether and to what extent these traits are interrelated and whether one of them could causally engender the others are not known.

**METHODS:** In a genetically selected rat model of reduced corticosterone responsiveness to stress, we explored PTSD-related biobehavioral traits using *ex vivo* magnetic resonance imaging, cued fear conditioning, and polysomnographic recordings combined with *in vivo* photometric measurements.

**RESULTS:** We showed that genetic selection for blunted glucocorticoid responsiveness leads to a correlated multitrait response, including impaired fear extinction, small hippocampal volume, and REMS disturbances, supporting their interrelatedness. Fear extinction deficits and concomitant disruptions in REMS could be normalized through postextinction corticosterone administration, causally implicating glucocorticoid deficiency in two core PTSD-related risk factors and manifestations. Furthermore, reduced REMS was accompanied by higher norepinephrine levels in the hippocampal dentate gyrus that were also reverted by postextinction corticosterone treatment.

**CONCLUSIONS:** Our results indicate a predominant role for glucocorticoid deficiency over the contribution of reduced hippocampal volume in engendering both REMS alterations and associated deficits in fear extinction consolidation, and causally implicate blunted glucocorticoids in sustaining neurophysiological disturbances leading to fear extinction deficits.

## INTRODUCTION

Posttraumatic stress disorder (PTSD) is the most prevalent psychopathological consequence of exposure to life-threatening or traumatic events (1,2) leading to both significant impairments in patients’ daily functioning and a substantial economic burden to society (3). Despite substantial progress in the development of therapies (4,5), a large group of patients show only partial long-term therapeutic efficacy, and their symptoms often relapse later in life.

A crucial challenge in PTSD research is understanding the substantial interindividual differences in vulnerability to developing PTSD (5), as only approximately 25-35% of persons exposed to severe trauma develop PTSD (1,5). Identifying predisposing factors is a crucial step not only for detecting individuals at risk but also for uncovering the mechanisms that trigger PTSD following trauma exposure and eventually for developing more effective (preventive or restoring) treatments (5).

Several behavioral (i.e., impairments in fear extinction) and biological (i.e., blunted glucocorticoid levels; small hippocampal volume; sleep disturbances) attributes exhibited as an individual’s preexisting traits have been identified as potential vulnerability factors for developing PTSD following trauma exposure. Behaviorally, a difficulty in extinction learning is a crucial trait, not only as a predisposition factor to develop posttraumatic stress symptoms (6–8) but also as a core process (i.e., the inability to extinguish a fear conditioning trace associated with the trauma) underlying the range of PTSD’s clinical manifestations (6,10–15).

Two biological traits, low glucocorticoid levels (15–17) and a smaller hippocampal volume (18–20), have been frequently reported in PTSD patients following trauma exposure in cross-sectional studies and were initially suspected to be a consequence of trauma exposure (21,22); moreover, these traits have received particular attention as potential preexisting PTSD risk factors (18,20,23). However, tackling their causal role has been a challenging task due to the difficulties of both collecting biological measures before trauma exposure and having access to relevant animal models in which the causal role of these traits can be investigated (24). A few longitudinal studies that have related pretrauma cortisol to subsequent PTSD development have yielded promising data regarding the predictive value of reactivity measures to real-life stressors, supporting a link between blunted cortisol responsiveness and PTSD development (25,26), while remaining inconclusive for the predictive value of basal cortisol levels (27–29). Regarding the hippocampus, evidence in favor of the vulnerability hypothesis for those with *a priori* smaller hippocampal volume to develop PTSD remains mixed. Evidence from twin (18,30) and longitudinal (20,31–33) studies has been both supportive (18,20,32,33) and disproving (30,31). Given the involvement of both glucocorticoids (24) and the hippocampus (34–36) in fear extinction, the consideration of blunted glucocorticoid responsiveness and smaller hippocampal volume as risk factors is still warranted, particularly as these aspects could relate to a particular subset of patients. Notably, the respective contribution of each of these traits to PTSD vulnerability remains unaddressed (37). This is particularly relevant, as the hippocampus is both a key regulator of the hypothalamus-pituitary-adrenal (HPA) axis, affecting both basal and stimulated glucocorticoid levels (38), and under the regulation of glucocorticoids, with both excess and deficient levels resulting in reduced hippocampal volume (39,40). To our knowledge, no study to date has integrated findings of blunted glucocorticoid responsiveness with structural brain imaging to ascertain the respective contribution of these traits to PTSD-related alterations.

Recently, sleep disturbances, particularly disruptions in REMS, frequently observed in PTSD patients (41–43) have also been proposed as a potential preexisting factor defining PTSD susceptibility (44). Sleep is key to memory consolidation processes (45,46), where glucocorticoid actions may be critical contributors (47,48), and sleep quality, particularly in the aftermath of traumatic exposure, has been underlined as a potential risk factor for the development of PTSD (49).

Here, we address two interrelated questions: 1) whether genetic selection for blunted glucocorticoid responsiveness can lead to changes in hippocampal volume, fear extinction and sleep architecture and 2) whether a constitutive deficit in glucocorticoid responsiveness not only predicts but is also causally implicated in the expression of fear extinction deficits—dissociated from any comorbid alteration in hippocampal volume—and sleep alterations. To this end, we make use of a genetically selected rat model of reduced corticosterone (CORT) responsiveness to stress (50–53) whose selection avoids inbreeding following a scheme of polygenic inheritance in alignment with the polygenic nature of both variation in plasma cortisol levels (54,55) and genetic risk for PTSD (56) in humans.

## METHODS AND MATERIALS

Detailed description is provided in the Supplement.

### Statistics

All data are represented as the means ± standard errors. The normality of the data distribution was assessed using the Kolmogorov‒Smirnov test. Unpaired *t* tests were used when only two groups were involved. When more than three groups were compared, one-way analysis of variance (ANOVA), followed by Holm‒Sidak *post hoc* tests, was used. When data were not normally distributed, Mann‒Whitney or Kruskal‒Wallis tests were used. Fear conditioning and *ex vivo* MRI datasets were analyzed using two-way ANOVA on repeated measures. Cumulative distributions were compared with the Kolmogorov‒Smirnov test. Data analysis was performed with GraphPad Prism 9 software using a critical probability of <0.05. Each datapoint in graphs represents an individual animal.

### Study approval

All procedures were conducted in accordance with the Swiss National Institutional Guidelines on Animal Experimentation and approved by the Swiss Cantonal Veterinary Office Committee for Animal Experimentation.

### Data availability

All data reported in this paper will be shared by the lead contact upon request.

## RESULTS

### Low glucocorticoid responsiveness to stress leads to a smaller hippocampus

To investigate whether constitutively low glucocorticoid responsiveness to stress leads to biobehavioral features involved in vulnerability to PTSD, we used three genetically selected rat lines (50–53) generated from an outbred population of Wistar Han rats and classified as Low-CORT responders (Low-CR), controls with normative CORT responsiveness (Ctr), and High-CORT responders (High-CR), (P30; Figure 1A and Figure S1A; see (53) and Supplement for details). We study the progeny of selected breeders (i.e., not exposed themselves to early-life stress) and specifically focus on the comparison between Low-CR and Ctr subjects. Accordingly, at adulthood, Low-CR rats exhibit blunted CORT responsiveness when exposed to novelty (open field; Figure 1B) or restraint (Figure S1B) stress. In addition, Low-CR display a blunted CORT circadian pattern in contrast to Ctrs, whose CORT levels increase toward the end of the light period (ZT 8-12) (Figure S1C).

**Figure 1.**
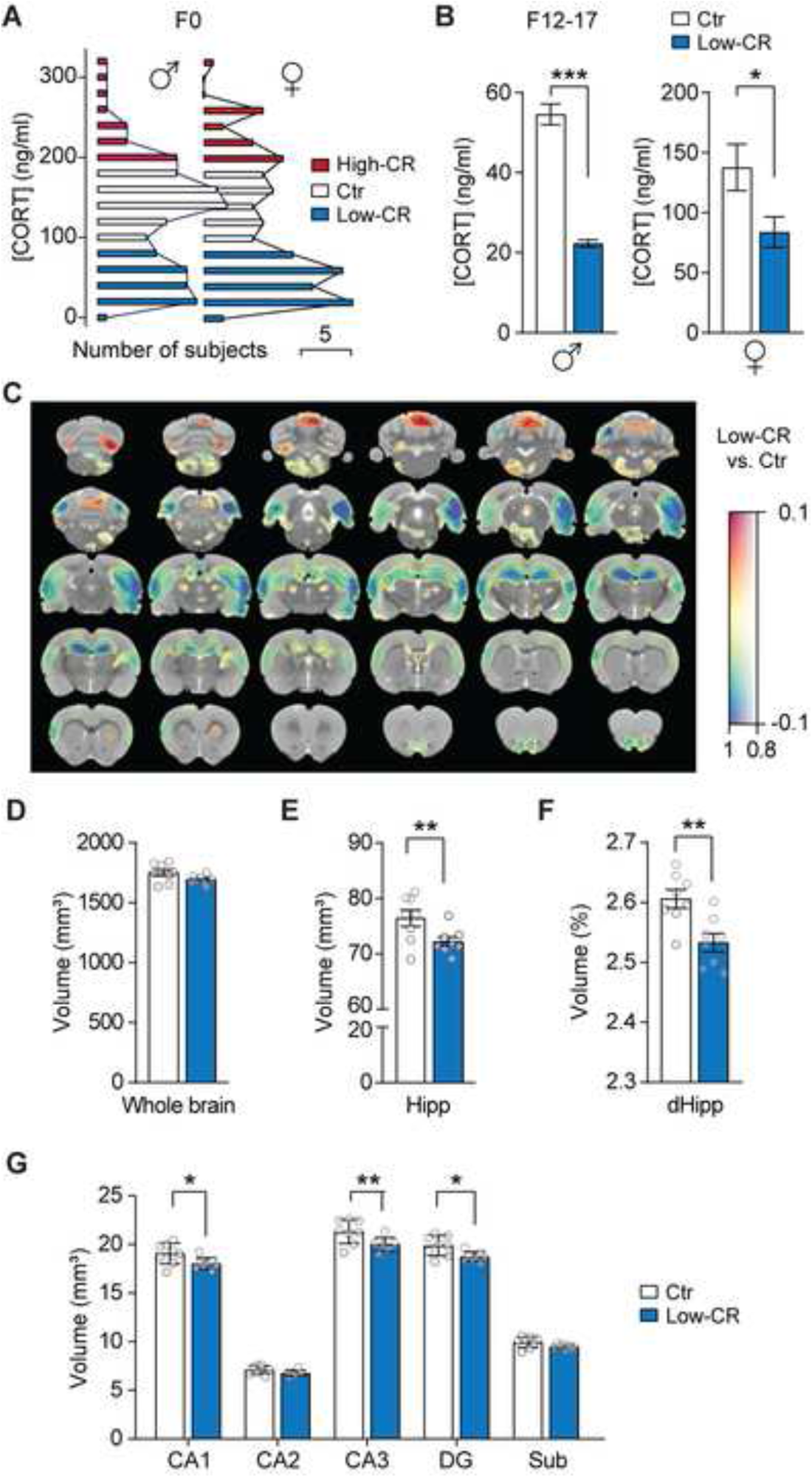
Constitutive differences in glucocorticoid responsiveness to stress are associated with hippocampal structural changes. (**A**) Distribution of plasma corticosterone (CORT) levels in response to stress exposure at the peripubertal stage in outbred Wistar male and female rats in the F0 generation reported in (53). Breeders were selected for the generation of lines with constitutive differences in glucocorticoid responsiveness, considering an upper limit of 100 ng/ml for Low-CR rats and a lower limit of 200 ng/ml for High-CR rats (color-coded). (**B**) Plasma CORT levels measured in the offspring from Ctr and Low-CR rats in adulthood after exposure to an open field for generations F12-F14-F17 (males; Ctr, n = 155; Low-CR, n = 238) and F13 (females; Ctr, n = 14; Low-CR, n = 12). (**C**) A map of voxelwise differences in GMV between Low-CR and Ctr rats calculated from *ex vivo* MR images overlaid on the rat brain MRI template. The colors of the overlay indicate brain volume differences (in mm^3^) thresholded at 1-p>0.8 (cool colors represent reduced volume, and warm colors represent increased volume in Low-CR compared to that in Ctr rats) and the transparency of the overlay is related to *P* value; black contours demarcate regions where uncorrected \**P* < 0.01. (**D** and **E**) Comparison of the volume of the total brain and of the whole hippocampus. (**F**) Comparison of dorsal hippocampus volume normalized to the whole brain. (**G**) Volumetric analysis of hippocampal subregions. Data are means ± SEMs; the number of observations is indicated by datapoints in the bar graphs. Statistical significance was assessed by the Mann‒Whitney test (B), unpaired *t* test (D-E-F), or two-way ANOVA (G), with \**P* < 0.05, \*\**P* < 0.01, \*\*\**P* < 0.001. Hipp, hippocampus; dHipp, dorsal hippocampus; CA, cornu ammonis; DG, dentate gyrus; Sub, subiculum.

First, we sought to determine whether Low-CR rats display differences in gray matter volume (GMV) across the brain. E*x vivo* MRI whole-brain *voxelwise* measurements (Figure 1C) indicated that while there were no differences in total brain size (Figure 1D), hippocampal areas are prominently decreased in Low-CR rats as compared to Ctr rats (Figure 1C). ROIwise analyses confirmed that Low-CR rats had smaller hippocampal volume than Ctr rats (Figure 1E, and Figure 1F with normalization to total brain volume), including differences in CA1, CA3, and DG (Figure 1G). To investigate the specificity of these findings, we measured a broad range of brain regions implicated in fear conditioning and extinction (57), including hippocampus, medial prefrontal cortex (mPFC), amygdala (Amy), and insular cortices. This analysis again highlighted a smaller volume in Low-CR rats circumscribed to the dorsal part of the hippocampus (Figure S1D).

### Low glucocorticoid responsiveness to stress leads to impaired fear extinction

Next, we asked whether blunted CORT responsiveness led to core behavioral features of PTSD, such as alterations in fear acquisition and/or extinction (5). We trained animals in an acoustic fear conditioning paradigm and then assessed their fear extinction capacities through exposure to repetitive CS presentations without US in a different context (Figure 2A).

**Figure 2.**
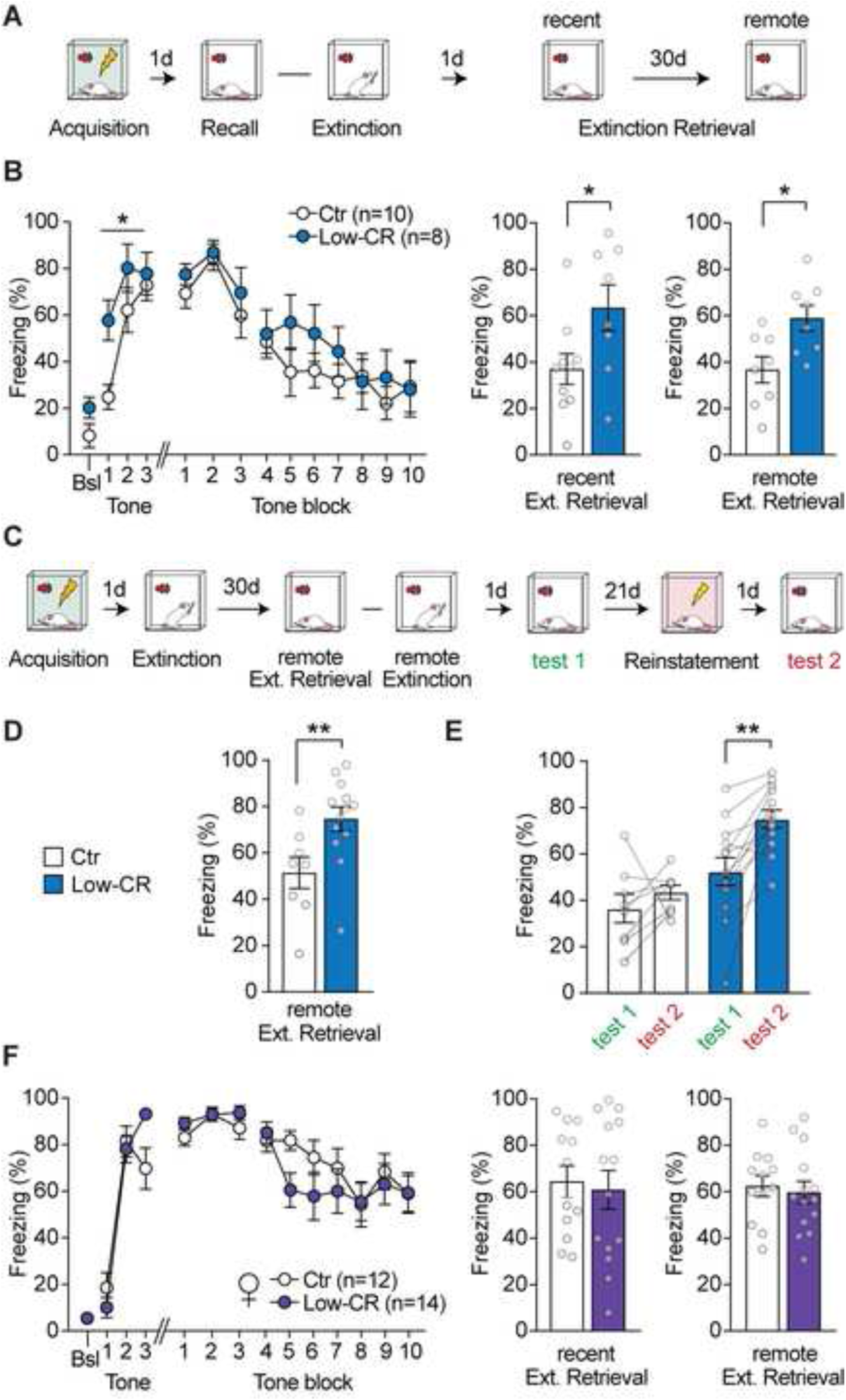
Blunted glucocorticoid responsiveness to stress is associated with impaired fear extinction in male rats. (**A**) Schematic representation of the paradigm for cued fear conditioning and extinction. The cued fear condition was induced by pairing a conditioned stimulus (CS, tone) with an unconditioned stimulus (US, foot-shock) three times. Memory recall and extinction training and retrieval were performed in a different context. Consolidation of extinction was assessed 1 day after extinction training (recent extinction retrieval) and 30 days thereafter (remote extinction retrieval). (**B**) Freezing levels measured in Ctr and Low-CR male rats during baseline (Bsl) and during the presentation of the CS in the acquisition phase, average freezing levels over 2 CS presentations (tone block) for recall and extinction training, and average freezing levels at recent and remote extinction retrieval. (**C**) Schematic representation of the paradigm for cued fear conditioning, extinction and subsequent fear reinstatement. (**D**) Mean freezing levels during remote extinction retrieval of rats subsequently exposed to fear reinstatement. (**E**) Fear relapse was assessed by comparison between the freezing levels during the last exposure to CS after remote extinction (test 1) and the freezing levels at CS presentation (test 2) after a mild shock was delivered in a novel context. (**F**) Same representation as in B for data from Ctr and Low-CR female rats. All numerical data are means ± SEMs; the number of observations is indicated by datapoints in the bar graphs. Statistical significance was assessed by unpaired *t* test (B-D-F) or RM two-way ANOVA (B-E-F) following Holm–Sidak post hoc analysis, with \**P* < 0.05, \*\**P* < 0.01, \*\*\**P* < 0.001. Nonsignificant comparisons are not indicated.

We started testing males and found that Low-CR do not differ from Ctr in the memory recall test given 24 h after training (Figure 2B) or during the subsequent extinction training (Figure 2B). However, Low-CR rats exhibited higher freezing levels than Ctr rats in the extinction retrieval session the next day. Their heightened fear response was still observed when testing took place 30 days later (remote extinction retrieval assessment; Figure 2B, D). These results indicate that Low-CR male rats show a deficit in the consolidation of fear extinction, a trait that predisposes humans to develop PTSD (6,7,58) and a distinctive hallmark of the disease (9–14). We replicated these extinction memory deficits in another cohort of animals (Figure 2C, D) that were further exposed to a reinstatement protocol (i.e., single US exposure in a novel context) 3 weeks afterward. Notably, Low-CR rats showed higher susceptibility to fear relapse, as indicated by their higher freezing levels to the CS in a recall session 24 h later (Figure 2D).

When the same tests were performed in females, we found no difference between Low-CR and Ctr rats in any of the fear processes examined (Figure 2E, F). Although we cannot exclude that Low-CR female rats would show fear extinction deficits if different training protocols were used, to advance the mechanistic understanding of the findings obtained in males, the remainder of the study was conducted in male subjects.

### Postextinction CORT treatment facilitates fear extinction memory in Low-CR rats

To probe the causal implication of blunted CORT responsiveness in the fear extinction deficits observed in male rats, we first confirmed that Low-CR displayed lower postextinction plasma CORT levels (Figure 3A, B). Then, we injected Low-CR with either CORT (1 mg/kg, i.p.) or vehicle immediately after extinction training. Extinction retrieval was tested in two sessions, i.e., on days 3 and 5 postextinction acquisition given that memory effects of exogenous glucocorticoids can take up to 48 hours after administration to emerge (59). Notably, whereas vehicle-injected Low-CR rats retained higher freezing levels than Ctr rats during the second extinction retrieval session, Low-CR rats injected with CORT showed a significant attenuation of the fear response (Figure 3C, Recent Ext. retrieval). This attenuation was long-lasting, as their freezing levels were still lower than in vehicle-injected Low-CR rats when at remote testing 30 days after fear acquisition (Figure 3C, Remote Ext. Retrieval). Furthermore, to assess the efficiency of corticosterone to continue supporting the extinction processes, Low-CR animals received a second CORT injection following a remote extinction session on day 30 (Figure 3C). Compared to vehicle-injected Low-CR rats, CORT-injected Low-CR rats showed attenuated freezing at remote extinction retrieval on day 33 (Figure S2A) and at fear relapse after reinstatement 3 weeks afterwards (Figure 3D). Taken together, these results indicate that postextinction CORT ‘normalizes’ fear extinction consolidation in Low-CR rats. These effects were long-lasting and specific to extinction memory, as the initial fear memory was still amenable to reinstatement.

**Figure 3.**
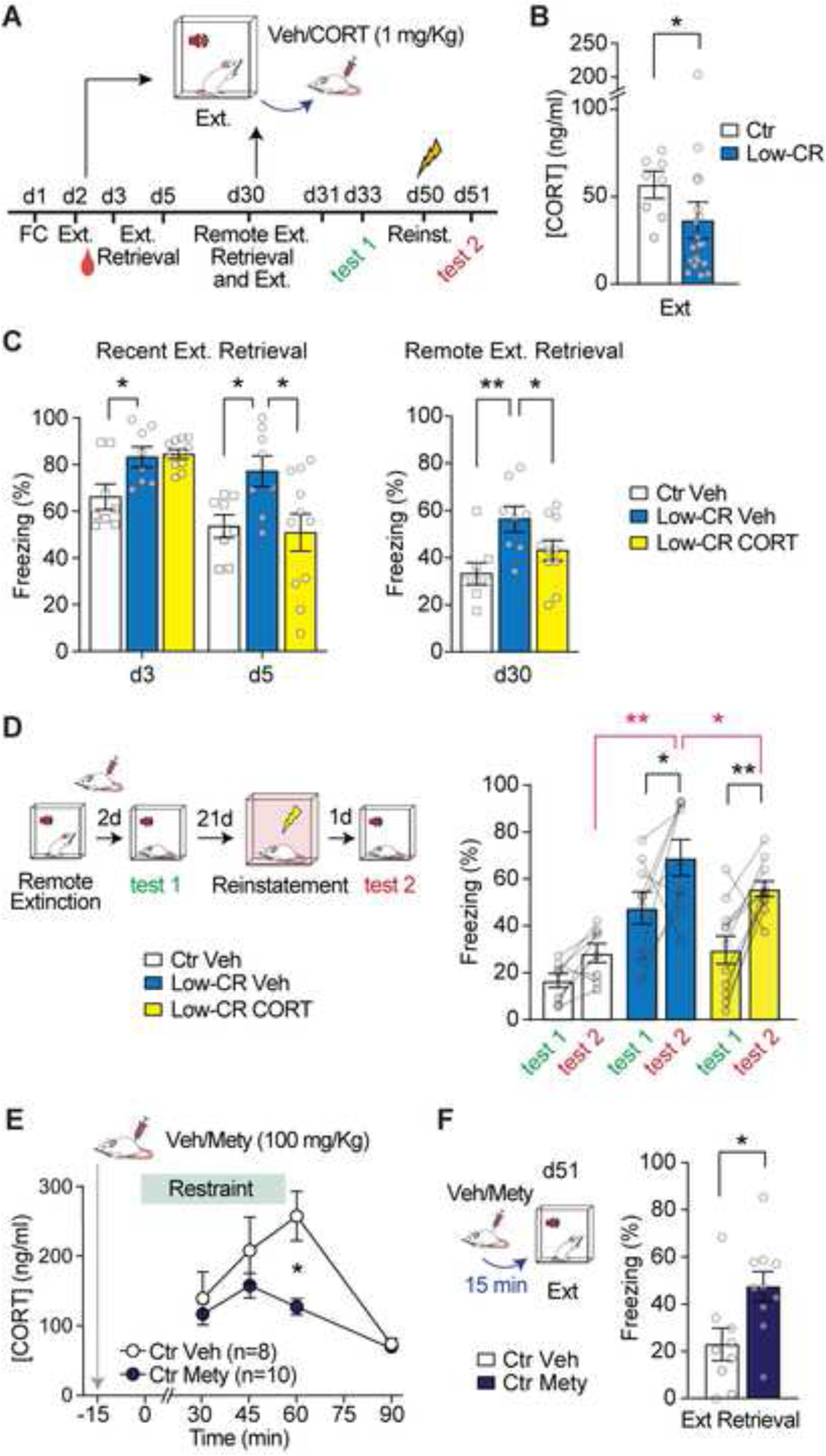
Modulation of glucocorticoids impacts fear extinction consolidation. (**A**) Schematic representation of the paradigm for cued fear conditioning and extinction with postextinction administration of CORT in Low-CR rats. (**B**) Plasma CORT levels measured in Ctr and Low-CR rats after the first extinction training. (**C**) Mean freezing levels during recent extinction retrieval (d3 and d5) and remote extinction retrieval (d30). (**D**) Mean freezing levels prior to (test 1) and after (test 2) fear reinstatement. (**E**) Time course of plasma levels of CORT in Ctr rats receiving an injection of vehicle/metyrapone (100 mg/kg) 15 min prior restraint. (**F**) Mean freezing levels during extinction retrieval in Ctr rats injected with vehicle or metyrapone before extinction training. All numerical data are means ± SEMs; the number of observations is indicated by datapoints in the bar graphs. Statistical significance was assessed by unpaired *t* test (F), Mann‒Whitney test (B), one-way (C) or RM two-way (D-E) ANOVA followed by Holm‒Sidak’s post hoc analysis, with \**P* < 0.05, \*\**P* < 0.01, \*\*\**P* < 0.001.

### Inhibiting CORT synthesis during extinction memory consolidation impairs extinction retrieval

As reported above, Low-CR rats present both blunted CORT responsiveness and reduced hippocampal size. To investigate whether reduced glucocorticoid release impacts fear extinction in isolation from any neuroanatomical difference, we assessed the impact of CORT synthesis inhibition in Ctr rats during the consolidation of fear extinction. We first verified the efficiency and time course of metyrapone (Mety; 100 mg/kg, s.c., 15 min prestress), a CORT synthesis blocker (60,61), in decreasing the levels of circulating CORT following exposure to restraint stress. Sixty min after stress onset, Mety-treated animals displayed blunted CORT levels compared to vehicle-treated animals (Figure 3E). Accordingly, we administered Mety 15 min before extinction training to dampen the levels of circulating CORT during the postextinction period. Freezing levels during extinction training were not affected by Mety (Figure S2B). However, in the extinction retrieval test administered 24 h later, Mety-injected animals showed higher freezing levels than vehicle-injected rats (Figure 3F). This indicated that decreasing CORT levels in the postextinction period inhibits the consolidation of extinction memory. When Mety was administered to affect CORT levels during the extinction training, no deficits at extinction retrieval were found (Figure S2C).

Together, these data indicate that CORT facilitates fear extinction memory and underscore the postextinction period as a critical time point for CORT actions, regardless of individuals’ hippocampal size.

### High glucocorticoid responsiveness is not associated with fear extinction deficits

Given that both blunted and high glucocorticoid levels can lead to reduced hippocampal volume (39,40), we made use of the High-CR line to assess how this trait relates to both hippocampal size and fear extinction. Notably, GMV analyses revealed that High-CR display smaller hippocampal sizes (Figure S3A, B C, E) with reductions restricted to the CA3 region (Figure S3D). Analyses including the fear conditioning and extinction network of brain regions did not identify any significant difference in other brain areas (Figure S3F).

Importantly, when exposed to a cue fear conditioning and extinction paradigm (Figure 4A), and despite their reduced hippocampal size, High-CR rats showed non significantly different freezing levels at extinction retrieval, spontaneous recovery (Figure 4B) and fear relapse (Figure 4C) than Ctr rats.

**Figure 4.**
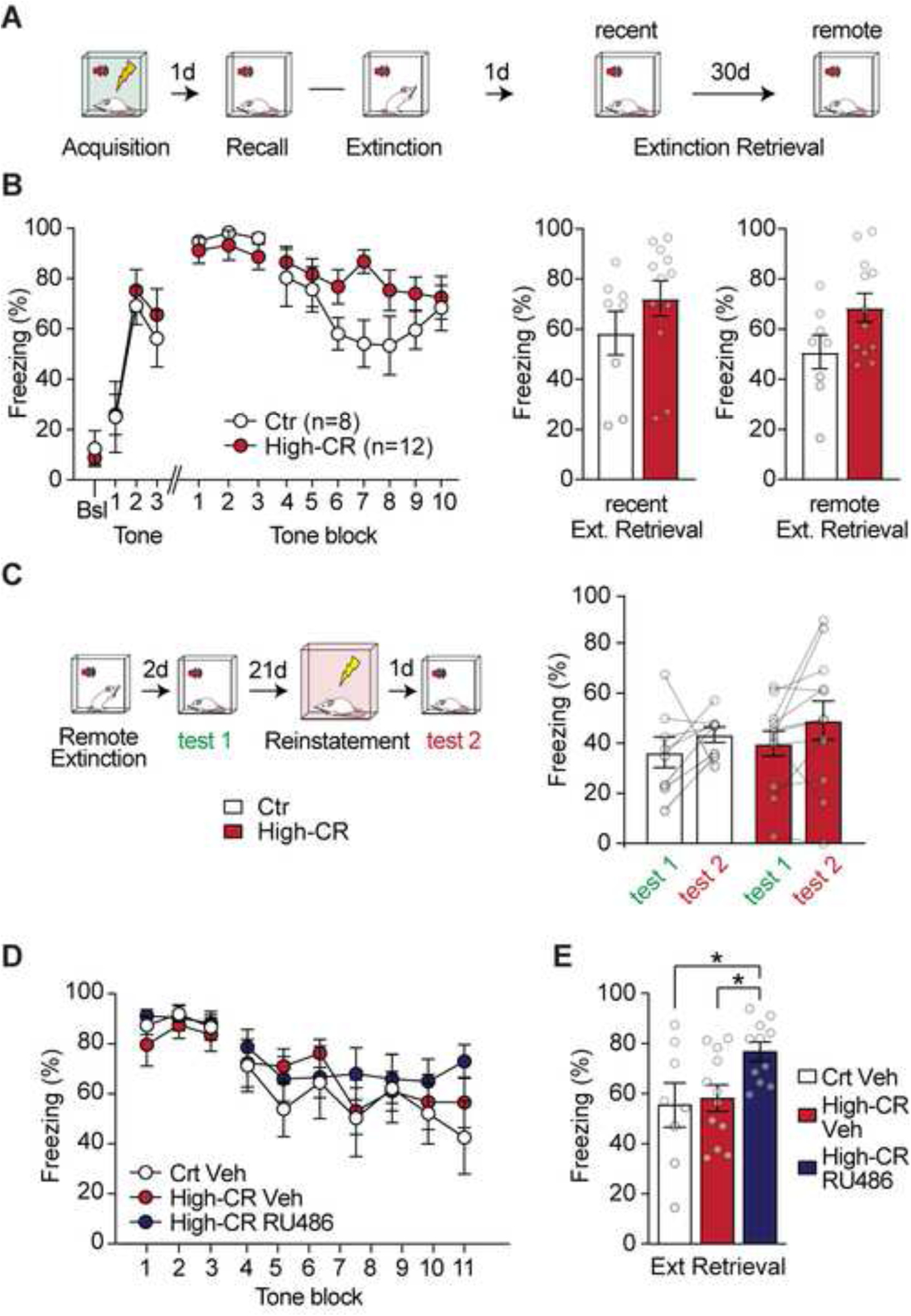
Elevated glucocorticoid responsiveness to stress is not associated with fear extinction impairments. (**A**) Schematic representation of the paradigm for cued fear conditioning and extinction. (**B**) Freezing levels measured in Ctr and High-CR male rats during baseline (Bsl) and during the presentation of the CS for the acquisition phase and average freezing levels over 2 CS presentations (tone block) for the postacquisition phases. (**C**) Schematic representation of the paradigm for fear reinstatement and mean freezing levels at subsequent recall for Ctr and High-CR rats. (**D**) Schematic representation of the paradigm for GR blockade using preextinction administration of RU486 (10 mg/kg) and freezing levels measured in Ctr and High-CR rats subsequently exposed to fear recall and extinction training. (**E**) Mean freezing levels during extinction retrieval in Ctr and High-CR rats injected with vehicle and High-CR rats injected with RU486 before extinction training. All numerical data are means ± SEMs; the number of observations is indicated by datapoints in the bar graphs. Statistical significance was assessed by unpaired *t* test (B), one-(E) or RM two-way (C) ANOVA followed by Holm‒ Sidak’s multiple comparison test, with \**P* < 0.05. Nonsignificant comparisons are not indicated.

To further investigate the relationship between hippocampal size and glucocorticoid actions on fear extinction, we next assessed the effects of glucocorticoid receptor (GR) blockade with RU486 (10 mg/kg; i.p.) in High-CR rats given just before the extinction session. While this treatment did not affect freezing during extinction training (Figure 4D), RU486-injected High-CR rats exhibited higher freezing responses than vehicle-treated High-CR and Ctr rats in the extinction retrieval test taking place 24 h afterwards (Figure 4E).

Together, these findings strongly support the view that blunted CORT responsiveness during the consolidation of fear extinction acts as a decisive factor in promoting fear extinction deficits, while small hippocampal size per se is not sufficient to determine impairments in fear extinction.

### Sleep-related signatures associated with blunted CORT responsiveness

Given the emerging hypotheses linking sleep disturbances with PTSD susceptibility (44,62,63), we performed polysomnographic recordings in naive Low-CR and Ctr rats during 48 h of undisturbed sleep/wake cycles (Figure 5A). Low-CR rats spent less time in REMS during the light (inactive) phase (Figure 5B) and more time in the awake state (Figure S4C), without differences in the number of bouts of the different sleep states (Figure S4A, B) or in the time in NREMS (Figure S4D). Notably, REMS bouts were longer in Low-CR rats (Figure 5C).

**Figure 5.**
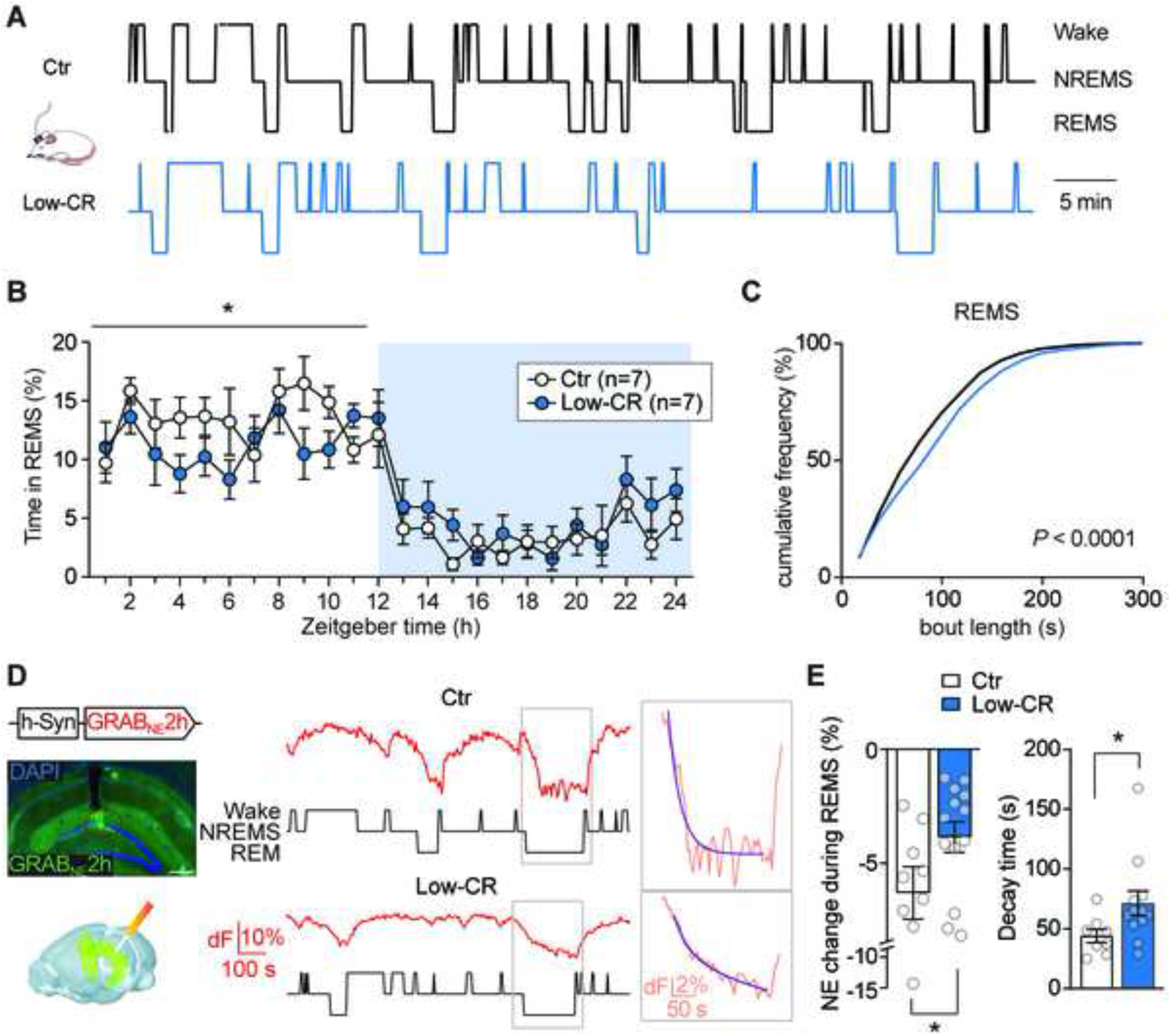
Blunted glucocorticoid responsiveness to stress is associated with REMS alterations. (**A**) Example hypnograms from a Ctr rat and a Low-CR rat that had received implants for EEG/EMG recordings. (**B**) Percent time Ctr and Low-CR rats spent in REMS across the sleep-wake cycle. (**C**) Cumulative distribution of REMS bout lengths across the sleep-wake cycle. (**D**) Schematic representation of photometric recordings in the hippocampus using the NE sensor GRAB_NE_2h, with the micrograph showing an example *post hoc* verification of the viral infection and recording site. Example traces show hippocampal NE levels (red) measured in a Ctr rat and a Low-CR rat and corresponding hypnogram (black), with enlarged portions showing biexponential fit (purple) of the photometric signal during REM episodes. (**E**) Mean values of the change in NE levels and decay times of NE signals (weighted τ from biexponential fit) measured during REMS in Ctr and Low-CR rats. The numerical data in B and E are means ± SEMs; the number of observations is indicated by datapoints in the bar graphs. Statistical significance was assessed by unpaired *t* test (E), Kolmogorov‒Smirnov test (C), RM two-way ANOVA (B), with \**P* < 0.05. Nonsignificant comparisons are not indicated.

(Norepinephrine) NE is emerging as a key candidate in linking the stress response, memory processing and sleep physiology (64–68). Excessive locus coeruleus (LC) activity and higher NE levels are reported in PTSD patients (69,70), and, in particular, heightened hippocampal NE signaling during REMS has been proposed to lead to PTSD pathophysiology (71). To investigate whether the altered REMS microarchitecture in Low-CR rats is associated with alterations in NE signaling, we combined polysomnographic recordings with fiber-photometry measurements using the fluorescent biosensor GRAB_NE_2h (65,68,72), which was expressed in the DG (Figure 5D). At NREMS-REMS transitions, we observed a pronounced drop in hippocampal NE in Ctr rats, consistent with the reported LC silencing during REMS (65,68,73). However, the change in NE levels was attenuated in Low-CR rats and was associated with a prolonged decay time compared to that of Ctr rats (Figure 5E), suggesting that a residual level of NE may persist in the DG during REMS. We found no differences in LC area, tyrosine hydroxylase (TH) density, or cFos activity in the LC between Ctr and Low-CR rats (Figure S5), suggesting that the differences in GRAB_NE_2h signal at NREM-REM transitions are not due to lower initial levels of NE release during NREMS in Low-CR rats, but rather to a persistence of extracellular NE during REMS.

Together, subjects with low CORT responsiveness exhibit a constitutive alteration in sleep architecture involving lower amounts of REMS, possibly accompanied by excessive NE signaling in the hippocampus.

### CORT normalizes REMS and hippocampal NE dynamics

Finally, we analyzed the sleep profile during the hours following extinction training (performed 5-6 h into the light phase), i.e., in the period expected to be conducive for the consolidation of fear extinction memory (74,75). Importantly, implanted Low-CR rats retained higher freezing levels than Ctr rats in response to the CS during the extinction retrieval test (Figure 6A).

**Figure 6.**
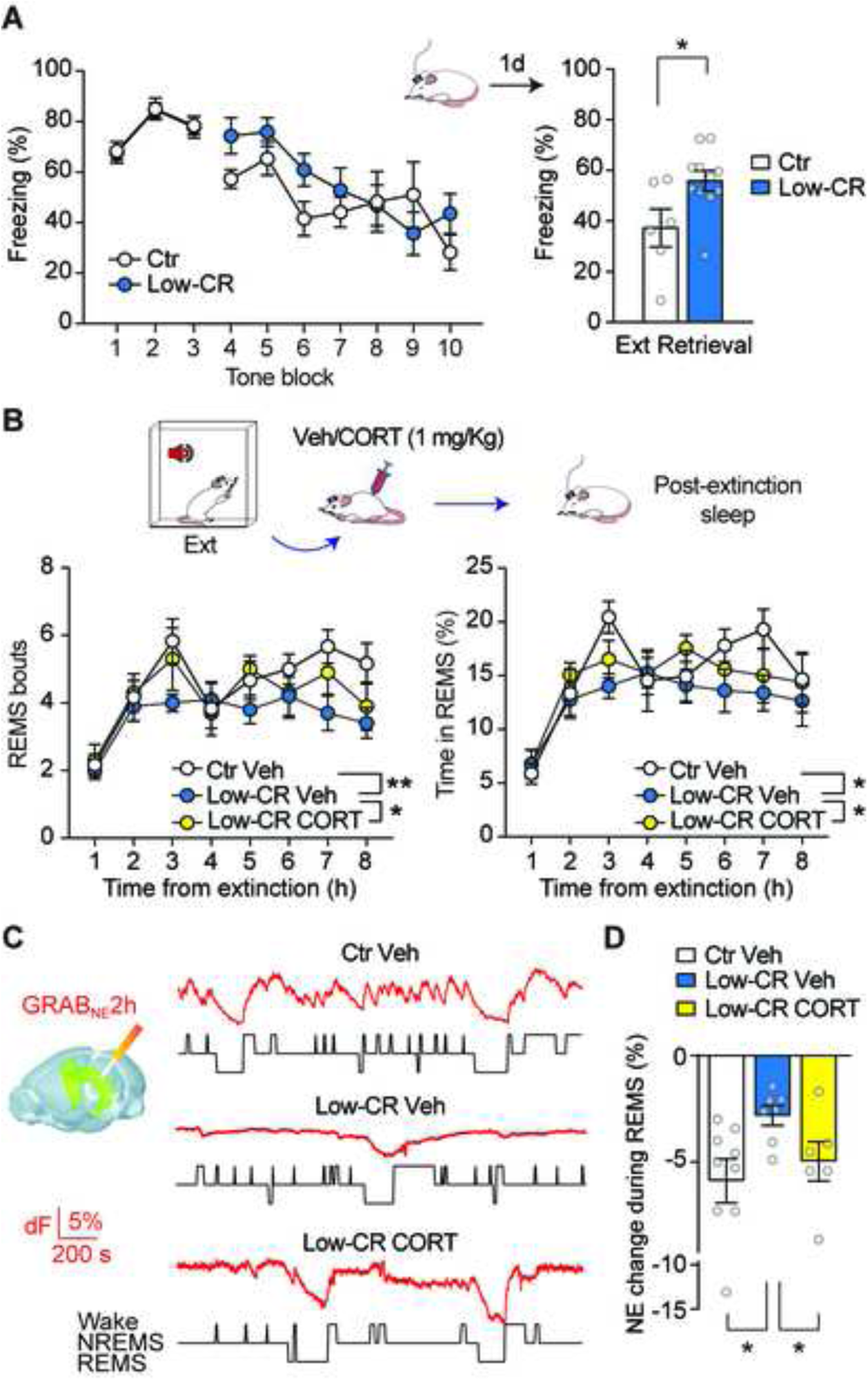
Postextinction REMS alterations are normalized by CORT administration. (**A**) Freezing levels during extinction training and extinction retrieval of Ctr and Low-CR rats that had received implants for EEG/EMG recordings. (**B**) Density of REMS bouts and the percent of time spent in REMS in the postextinction period for Ctr and Low-CR rats injected with vehicle and Low-CR rats injected with CORT (1 mg/kg) after the extinction training session. (**C**) Schematic representation of photometric recordings in the hippocampus using the NE sensor GRAB_NE_2h, with example NE signals (red) and corresponding hypnograms (black) measured in a vehicle-treated Ctr and Low-CR rat and a Low-CR rat injected with CORT. (**D**) Mean values of the change in NE levels measured during REMS in Ctr and Low-CR rats. The numerical data are means ± SEMs; the number of observations is indicated by datapoints in the bar graphs. Statistical significance was assessed by unpaired *t* test (A, right), Mann‒Whitney test (D), RM two-way ANOVA (A, left-B), with \**P* < 0.05, \*\**P* < 0.01. Nonsignificant comparisons are not indicated.

The hippocampus contributes to long-term consolidation during sleep, even for nonclassical hippocampus-dependent memories (76,77). Thus, we examined REMS architecture and hippocampal NE levels in CORT-injected Low-CR rats and vehicle-injected Low-CR and Ctr rats. The CORT injection (1 mg/kg, s.c.) was administered immediately after extinction training, after which rats were tethered to the EEG/EMG system and left undisturbed in their home cage. Ctr rats showed a surge in REMS in the first three hours postextinction, which appeared as an increase in both the number of REMS bouts and the percent of time spent in REMS, a phenomenon that has been reported to support the consolidation of fear extinction (45,74,75,78)(Figure 6B). However, vehicle-treated Low-CR rats failed to show this surge in REMS, and both the bout number and the percent of time spent in REMS reached a plateau once sleep was stabilized after the first hour postextinction training. Notably, CORT treatment in Low-CR rats significantly affected these parameters, which were normalized toward levels found in Ctr rats.

Photometric recordings in the hippocampus revealed that the NE levels during REMS in Low-CR rats remained more elevated than those in Ctr rats during the postextinction training period (Figure 6C, D), expanding our baseline observations (Figure 5E). Importantly, CORT treatment reduced NE levels during REMS leading to a normalization of the excess in hippocampal NE observed in Low-CR rats.

Together, these data underscore the relevance of REMS during the postextinction training phase for the consolidation of extinction memory and support the implication of blunted CORT responsiveness in the alterations of REMS architecture and hippocampal NE signaling that are detrimental for memory consolidation.

## DISCUSSION

We show here that genetic selection for blunted glucocorticoid responsiveness (50–53) engenders a distinct phenotype encompassing several pivotal PTSD vulnerability traits, including impaired fear extinction, small hippocampal volume and REMS disturbances, revealing that these traits are biologically interconnected. This is quite remarkable, as most studies typically focus on one (maximum two) risk factor(s) at a time, considering them as independent risk factors. We demonstrate a causal role for a deficiency in mounting a glucocorticoid response in both fear extinction deficits and concomitant REMS disruptions, which were normalized by CORT administration during the postextinction period.

PTSD patients frequently show smaller hippocampal volume. Several human studies have proposed that a reduced hippocampal volume may constitute a vulnerability factor for PTSD development (18,20,79), while others did not find support for this view (31) suggesting that the reduction frequently observed in PTSD individuals is mainly due to environmental effects, such as the stress of combat (30). Our study provides strong data in support of a preponderant role for glucocorticoid deficiency in engendering both REMS alterations and associated deficits in fear extinction consolidation while minimizing the contribution of reduced hippocampal volume *per se*. The reduction in hippocampal volume observed in Low-CR rats is consistent with evidence reporting a bidirectional relationship between glucocorticoid levels and hippocampal structure (38–40). Given that the reductions we observed were in animals not exposed to stress or trauma, they bring forward the difference in hippocampal size as a further constitutive trait to consider in interpreting our data.

Low-CR rats show a flattened circadian pattern but, importantly, it was sufficient to provide a corticosterone treatment during the fear extinction consolidation period ‘to correct’ their impaired fear conditioning. These data help explaining clinical data indicating that, whereas blunted cortisol from pretrauma reactivity measures to life stressors predict PTSD development well (25,26), basal cortisol levels are not good predictors (27–29). Fear extinction deficits in PTSD patients are typically not manifested during extinction training but are particularly observed in subsequent extinction recall sessions (12,58,80), pointing to a specific deficit in the ‘consolidation’ of fear extinction. This link between blunted corticosterone responsiveness and poor extinction phenotypes aligns well with the well-known facilitatory role of glucocorticoids on memory consolidation (81–83), including fear extinction memory (84–87).

We took advantage of the characteristics of all the CORT rat lines developed in the laboratory, including the High-CR line, to parametrically investigating how variations in glucocorticoid function during the postextinction period and hippocampal size relate to the efficiency of fear extinction consolidation. First, the data from the three rat lines indicated an inverted-U shape relationship between constitutional glucocorticoid responsiveness and hippocampal size, consistent with previous observations in humans (88). Then, we implemented gain- or loss-of-function pharmacological manipulations to modify CORT levels or their actions through GRs during the fear extinction consolidation period and found that i) as indicated above, in Low-CR rats, which showed impaired extinction and reduced hippocampal volume, increasing systemic CORT levels rescued extinction memory; ii) in Ctr rats, which showed normative levels of fear extinction, CORT responsiveness and hippocampal size, blocking CORT synthesis led to increased freezing levels at subsequent extinction retrieval; iii) in High-CR rats, which did not show impairments in fear extinction or fear relapse at reinstatement but displayed reduced hippocampal volume, antagonizing GRs impaired the consolidation of fear extinction. Overall, these observations indicate that blunted, but not high, glucocorticoid responsiveness is associated with impaired fear extinction memory consolidation. These findings also strongly implicated CORT signaling in proper consolidation of the extinction memory and suggest that hippocampal volume may not relate to fear extinction deficits per se but through its capacity to regulate HPA axis functioning and glucocorticoid signaling (37,38).

Sleep disturbances, in particular REMS, are not only a major symptom of PSTD (41–43) linked to negative outcomes (89,90), they have also been proposed to play an etiological role in PTSD (44) and predict worse disease symptomatology following trauma exposure (62,63,91). In line with these data, Low-CR rats spent less time in REMS during their inactive phase. Postextinction REMS is required for the successful consolidation of the extinction memory (74,75). In our study, the failure of Low-CR rats to achieve sufficient REMS after extinction aligns with data showing that REMS deprivation immediately after extinction training led to fear extinction deficits in rats (74).

During REMS, the brain experiences a characteristic neuromodulatory profile involving high glucocorticoid and low catecholamine levels, including NE (46). The LC decreases its activity and shuts down immediately before and during REMS (73), and these silent LC periods during REMS are crucial for memory consolidation (64,68). Heightened LC-NE activity is also a recognized alteration associated with PTSD and has been implicated in hyperarousal during sleep (69,70,92). Our observations suggest that Low-CR rats exhibit excessive hippocampal NE levels during REMS. These findings provide the first experimental evidence in support of a recent hypothesis positing that a high NE tone during REMS is causally involved in the failure to consolidate processes of the extinction memory (71). Ultimately, during NE-enriched REMS, also referred to as “restless REMS” (93), a failure of the hippocampus-PFC-Amy network is responsible for an impaired consolidation of extinction memory.

Therefore, in our study, REMS arises as an important physiological factor that is highly modulated by variations in glucocorticoid responsiveness and, in parallel, is strongly involved in the adaptive consolidation of cued fear extinction memory.

Our data identify the postextinction training period as the crucial time point at which a glucocorticoid deficiency should be targeted with exogenous treatments and identify the restoration of sleep architecture as a likely mechanism of action implicated in glucocorticoid effects. Several studies have shown that glucocorticoid administration facilitates fear extinction (85,94–97). However, only a limited number of studies have tried this intervention in PTSD patients (98). Our findings underscore the importance of considering interindividual differences in HPA axis responsiveness when evaluating a clinical strategy and highlight the postextinction phase as a critical moment for glucocorticoid administration.

Strikingly, our genetic selection scheme, while being highly effective in attribute selection (i.e., achieving a specific range of CORT responsiveness within each line), concurrently leads to small hippocampal volume, impaired fear extinction, and REMS disturbances, each of them being important PTSD vulnerability traits. Unlike previous PTSD studies that focused on 1—or a maximum of 2—traits our genetic selection approach revealed the interconnectedness of these traits. The reasons for their interconnectivity can be multifactorial, including both genetic (e.g., pleiotropy, genetic correlation, and linkage disequilibrium) and nongenetic (e.g., mechanistic effects, where one trait affects the physiological or developmental processes that lead to changes in the other traits) factors. Glucocorticoids—our targeted trait for selection—through their genomic and nongenomic effects can regulate hippocampal structure, memory consolidation and sleep architecture (82,100–103). Our own findings support the causal role of glucocorticoids in extinction memory and REMS alterations. In turn, changes in some of these traits (e.g., sleep disturbances, particularly in REMS processes) can also influence the expression of other traits (e.g., the consolidation of extinction memory)(47,78).

Several reasons support the view that the multitrait interconnectedness found following selection for blunted glucocorticoid responsiveness is representative of their relationship in nature. First, our genetic selection scheme followed gold standard artificial selection approaches to maximize genetic variance in a unique population while facilitating selection response (104). The parental group was a large sample of outbred Wistar rats with minimum coancestry. Selection criteria over generations allocated animals to the different CORT groups according to fixed measures—as opposed to relative percentile cutoffs for each cohort—which ensured a robust selection response. Second, a crucial aspect of our nonrandom mating procedures for line generation (i.e., sibling mating was excluded, and a refreshing of the selection introduced in F12) was inbreeding avoidance which favors genetic diversity (105) and polygenic inheritance (106). The latter is crucial to align with the polygenic nature of both variation in plasma cortisol levels (54,55) and genetic risk for PTSD (56) in humans. Third, for genetic selection schemes that avoid inbreeding, as in our case, the risk for genetic drift -and associated instability of correlational traits-is minor. Finally, the effects of CORT treatment indicate a causal role of blunted glucocorticoid responsiveness on fear extinction deficits and REM alterations.

Our study had some limitations. It is somewhat surprising that we did not find fear extinction impairment in female Low-CR rats, given the reported higher prevalence of PTSD in women (107). However, the type of trauma and the age at the event affect PTSD development in a sex-dependent manner (1,108). Delahanty et al. (2005) showed a sex-specific relationship between increased peri-trauma cortisol levels and increased PTSD symptoms in male children but not female children. These observations suggest the presence of potential sex differences in biological predictors and PTSD vulnerability.

## AUTHOR CONTRIBUTIONS

CS, SA and SM conceived the study. SM, SA, IGdS, JG, OZ, SEW, MM, TCW and DC acquired and analyzed data. CS and DC acquired funding. CS, SA and SM wrote the manuscript. SA and SM prepared manuscript figures. All authors read, edited and approved the final manuscript.

## ACKNOWLEDGEMENTS

We thank the members of the Laboratory of Behavioral Genetics that contributed to the development and maintenance of the rat lines across generations, in particular Damien Huzard. We thank João Paulo Brás for performing behavioral experiments in female rats, Laurine Lang for developing the protocol for CORT extraction from fecal boli and Malina Vaucher, Estelle Renard-Dausset, Flora Maurici and Cristian Pérez-Fernández for histological analyses. The authors are grateful to Anita Lüthi for insightful discussion and critical input. This work was performed in part using the resources and services of the BIOP and CPG Research Core Facilities at the School of Life Sciences of EPFL.

This study has been supported by grants from the Swiss National Science Foundation (NCCR Synapsy, grant no. 51NF40-158776), European Union’s Seventh Framework Program for research, technological development and demonstration under grant agreement no. 603016 (MATRICS), ERA-NET Neuron (SNSF Project No. 31NE30_189061 Biostress) and intramural funding from the EPFL.

