## Supplemental Methods and Suppl Figures for "Blunted glucocorticoid responsiveness to stress causes behavioral and biological alterations that lead to posttraumatic stress disorder vulnerability"

**The document includes:**

Supplemental Methods

Supplemental References

Supplemental Figures S1 to S5

### Supplemental Methods

*Animals*

Experimental animals (males and females) were offspring from F8-F17 generations of genetically selected corticosterone stress lines of Wistar Han rats (1,2) developed and bred in our in-house animal facility (EPFL, Lausanne, Switzerland). Of note, experimental animals used in this study were not exposed to the selection procedure for CORT reactivity ("corticosterone-adaptation-stress-test", see below). Rats were maintained on a 12-h light–dark cycle (lights on at 07:00 h) in a temperature- (21 ± 1°C) and humidity-controlled environment (55 ± 5% humidity) with *ad libitum* access to laboratory chow and water. At weaning on postnatal day 21 (p21), rats from different litters were matched according to weight and grouped-housed by three animals per cage. Rats then remained undisturbed, except for weekly cage change and body weight monitoring at p28, p30, p36, p42, and p60, until experimental procedures began in adulthood (p90). Rats were handled for three consecutive days before the first behavioral test was performed. Experiments were performed between 07:30 h and 13:00 h unless stated otherwise. All procedures were conducted in accordance with the Swiss National Institutional Guidelines on Animal Experimentation and approved by a license from the Swiss Cantonal Veterinary Office Committee for Animal Experimentation.

*Protocol for selective breeding*

The selected rat lines used in this study were derived from outbred Wistar Han rats obtained from a commercial provider (Charles River, France) and bred over generations in our animal facility, as previously described (1,2). Briefly, for every generation, breeders were selected based on their CORT responsiveness following three days of repeated stressors at the prepubertal stage (p28-p30). The stressors included exposure to an open arena (5 min), to an elevated platform under bright light (EP; 25 min) and to a synthetic predator odor (trimethylthiazoline, TMT, Phero Tech Inc.; 25 min). At p28 and p30, blood was sampled from animals by means of a tail incision. For each breeding generation, selection was based on plasma CORT concentration in both males and females at the offset of a stress exposure period of 50 min to 2 consecutive stressors (TMT followed by EP) on p30. Specifically, animals were segregated as "Low-CORT responders" (Low-CR; CORT< 100 ng/ml), "Normative CORT responders" (i.e., controls or "Ctr"; CORT > 100 ng/ml and < 200 ng/ml), and "High-CORT responders" (High-CR; CORT > 200 ng/ml), and selective intraline breeding was carried out over several generations (see Figure 1A and Supplemental Figure 1A). In each generation, males and females served as founder pairs to generate the following generation and experimental animals. To minimize the effects of both extreme inbreeding and genetic drift, the breeding scheme strictly excluded sibling and backcrossing mating. Consequently, the breeding scheme followed a polygenic selection scheme (3). Importantly, experiments in this study were performed on the offspring of selected breeding animals (and, therefore, these animals were not exposed to prepubertal stress themselves), and the main focus of the study was on Low-CR and Ctr lines, although a few experiments were also performed in the High-CR line (c.f., specific protocols).

*Open field (OF)*

The OF test was used to assess reaction to novelty and exploratory behavior, as described previously (4). The OF consisted of a circular arena (1.50 m diameter, with a 32 cm high wall), with light adjusted to a level of 8-10 lx and 3-5 lx in the center and at the wall, respectively. Animals were introduced into the arena facing the wall and were allowed to freely explore for 20 min. The animals' positions were monitored using a video camera mounted above the center of the arena. For analysis, the arena was divided into three virtual concentric parts: a center, an intermediate, and a wall zone. A computerized tracking system (Ethovision 11.0 XT, Noldus, Information Technology) recorded the time spent exploring each zone as well as the distance moved. The apparatus was cleaned with a 5% ethanol solution between trials.

*Cued fear learning paradigms*

Fear conditioning and extinction took place in two different contexts (A and B). Additionally, a third context (C) was used during reinstatement. The training cage (Context A) consisted of two aluminum sidewalls, a Plexiglas ceiling, a white rear wall, and a Plexiglas door (30 × 24 × 21 cm; Med-Associates, St Albans, VT). The floor consisted of stainless-steel rods connected to a shock source (Med-Associates) for the delivery of foot shocks. Context B consisted of a white Teflon cylindrical chamber with a plastic flat floor. In addition, a paper cylinder and white paper were placed in front of the Plexiglas door. Context C was similar to Context A, except for a black rear wall and a different floor where steel rods were larger in size. The chambers were located inside a sound-attenuating box and were cleaned with 1% acetic acid (for Context A) or with 5% ethanol (for contexts B and C) before and after each session. Ventilation fans, a speaker, and house lights were also present in each chamber. Stimuli (light, paper cylinder or white paper, and odor) were adjusted within conditioning chambers to generate three distinct contexts. In all sessions, the animals' behaviors were monitored with a camera connected to a video recorder. Behavioral freezing, defined as the absence of all bodily movements except breathing-related movement (5), was used as the fear response measure. Freezing epochs, defined as a period of immobility of at least 1 s, were scored either manually by an individual blind to the experimental conditions or automatically through Video Freeze Software (Med-Associates).

<u>Conditioning</u>

Fear conditioning (FC) was performed in a single session in the afternoon. After 3 min of free exploration in Context A, rats received three pairings of a 20-s CS tone (80 dB, 820 Hz) ending with a 1-s US foot-shock (0.5 mA). The intershock interval was 60 s. After the last shock, the animals were kept in the conditioning chamber for an additional 30 s. The context was scented with citronella, with fans on, and exposed to 40 lx light. For the metyrapone experiment, slightly different conditioning was used: after baseline, 3 CSs (90 dB, 5 kHz, 25 s) were presented, ending with 1-s US of 0.5 mA intensity.

<u>Extinction training</u>

Extinction of cued fear learning took place one day after FC in the mornings in Context B, lit in green and scented with lavender, with fans off. Animals were free to explore the environment during the first 3 min, and then 20 CSs were presented every 60 s. Animals were kept 60 s after the last CS for a total duration of 30 min. A remote extinction was run in the same context after 30 days, which was mentioned in the results. During remote testing, freezing during the first six tones was considered a measure of spontaneous recovery (SR). For the metyrapone experiment, 24 CS (90 dB, 5 kHz, 25 s) were presented every 50 s after a 3 min baseline. In the text, the first period of exposure to 6 tones is referred to "recall" and the following period is referred to as "extinction".

<u>Extinction retrieval</u>

Twenty-four and forty-eight hours after training or twenty-four hours after reinstatement, extinction memory was assessed in Context B. After 3 min of free exploration, the rats experienced six CS presentations with an intertrial interval of 60 s. Rats were removed from the chambers 60 s after the final CS presentation (11 min total duration). The same procedure was followed at remote timing. For the metyrapone experiments, eight CSs were presented for a total time of 13 min.

Reinstatement

Three weeks after the last remote cue test, animals were placed in Context C for 3 min. After 120 s, a subthreshold foot-shock (0.4 mA) was delivered for 1 s. Animals were removed from the chambers 60 s after the shock. The context had a yellow light with the fan on and had a rose scent.

*Corticosterone measurement from blood samples*

Blood samples were obtained by tail-nick and collected into heparin-coated capillary tubes (Sarsted) that were kept on ice until centrifugation (4 min, 4°C and 10000 rpm). Plasma samples were stored at −20°C. Plasma CORT levels were measured using a highly sensitive ELISA kit (1/40 dilution, Enzo Life Sciences) according to the manufacturer's instructions. CORT levels were calculated using a standard curve method.

*Circadian corticosterone pattern measurement from fecal boli*

To assess the circadian corticosterone pattern in the Low-CORT and Ctr lines, rats were housed individually in cages with a grid floor allowing for collection of fecal boli from the bedding underneath. After three days of adaptation, samples were collected for 24 h at 4-h intervals in Whirl-Pak bags (Nasco, Fort Atkinson, WI, USA) and frozen at -20°C. Corticosterone extraction was performed as previously described (6), with some modifications. Briefly, frozen samples were crushed into powder, mixed with 95% ethanol (10 mL/g) in tubes, and incubated overnight at room temperature on a roller mixer. After centrifugation (20 min at 20°C, 600x g), 100 µL of the supernatant was transferred into fresh tubes that were placed on a heater (37°C) until the liquid evaporated completely. The solid residue was resuspended in 200 µL of assay buffer (Corticosterone ELISA kit, Enzo Life Sciences) and frozen at -20°C until use. CORT levels were measured using a highly sensitive ELISA kit (1/40 dilution, Enzo Life Sciences) according to the manufacturer's instructions. The obtained values of CORT were normalized to the weight of the fecal samples after they were dried at 37°C. For the representation of these data, CORT values measured from samples collected in a given 4-h interval are assigned to the Zeitgeber time corresponding to 9 h earlier (i.e., 5 h before the beginning of the 4-h interval), in order to consider the time shift between fecal CORT metabolites and plasma CORT levels (6).

*Drugs*

CORT (Sigma-Aldrich) was injected intraperitoneally at a dose of 1 mg/kg in a vehicle 0.9% NaCl. Glucocorticoid inhibition was induced by injecting either RU486 (mifepristone, Sigma-Aldrich), an antagonist of glucocorticoid receptors, or metyrapone (Tocris Bioscience), a blocker of CORT synthesis and release. RU486 was dissolved in saline and injected intraperitoneally (10 mg/kg) 30 min before extinction training. Metyrapone was dissolved in ethylene glycol and diluted in saline, and injected subcutaneously (50-100 mg/kg) either 90 or 15 min before extinction training.

*Brain fixation for anatomical studies*

Male animals from the F8 generation were used for anatomical studies. At p90, rats were anesthetized with a lethal dose of pentobarbital (intraperitoneally, 150 mg/kg; Esconarkon, Streuli Pharma) and transcardially perfused using 0.9% saline solution followed by a fixative solution of 4% paraformaldehyde in phosphate-buffered saline (PBS, pH 7.4). The animals were decapitated; then, the heads were stored in 4% paraformaldehyde overnight and rehydrated in PBS containing 0.05% sodium azide for at least one week before scanning.

*Tissue preparation and immunofluorescence*

Rats were deeply anesthetized with pentobarbital (150 mg/kg intraperitoneally, Streuli Pharma) and perfused transcardially (4% PFA, 1× PBS, pH 7.4). The brains were extracted and placed in 4% PFA at 4°C for 24 h, following incubation for 48h in a 30% sucrose solution at 4°C. Frozen brains were cut into 30 µm coronal sections with a cryostat (Leica, Wetzlar, Germany) and stored into a cryoprotectant medium. Free-floating sections were incubated in blocking solution were blocked in 1x PBS with 5% normal donkey serum (NDS, Jackson ImmunoResearch Laboratories, West Grove, PA, USA) and 0.3% Triton X-100 (Sigma-Aldrich) for 90 min at room temperature (RT), followed by overnight incubation in a freshly prepared antibody solution (1x PBS / 1% BSA / 0.3% Triton X-100) at 4 °C with rabbit anti-cFos (1:1000, rabbit anti-cFos, no. 226003, Synaptic System), and/or mouse anti-TH (1:2000; mouse anti-TH, Immunostar), under constant shaking. Sections were washed in 1x PBS and fluorophore-labeled anti-rabbit (1:500, AlexaFluor-488, Life Technologies), and/or anti-mouse (1:500, AlexaFluor-647) IgGs were used as secondary antibodies, in antibody solution for 2 h at RT. DAPI (1:10000 in 1x PBS) was used to stain the nuclei (10 min at RT). Sections were washed again with 1x PBS and mounted on Superfrost glass slides (Thermo Fisher Scientific, Waltham, MA, USA) with Fluoromount mounting medium (Southern Biotech, Birmingham, AL, USA).

For verification of viral infection with $GRAB_{NE}$2h and fiber optic placement, sections were incubated with goat anti-GFP (Abcam, 1:500), followed by anti-goat (AlexaFluor-568; 1:1000), and with DAPI.

*Imaging and quantification*

For brain activation study with anti-cFos and for viral infection verification with anti-GFP antibodies, images were acquired on a virtual slide microscope (VS120, Olympus) with an ×20 objective. For LC area measurement and tyrosine hydroxylase (TH) quantification, images were acquired on a Leica SP8 confocal microscope slide scanner using a 63x/1.40 oil objective. Regions of interest were manually outlined based on the Paxinos rat brain atlas. For cFos+, GFP+, and DAPI+ cell detection, images were analyzed with QuPath v0.1.4 and v.0.2.3, using a custom-built script. The density of cFos-positive cells (cFos+/$mm^2$) was averaged over 3-4 sections per animal. To quantify TH expression, the background of the channel was measured at three different random areas around the section and averaged together to generate a mean background which was then subtracted from the TH channel. Image analysis and cell counting were performed using ImageJ software. Sections were then averaged to provide one value per animal per cell type of interest. For viral infection, the signal from the fluorescent reporter was manually thresholded and quantified with QuPath. The DG area and its surrounding parts were carefully screened for possible off-site injection. Optic fiber placement was verified based on fiber tract lesion. Animals that showed viral expression and optic fiber outside the targeted area were excluded from the behavioral analysis.

*Ex-vivo MRI*

Imaging was conducted as previously described (7). Before scanning, the lower jaw was removed from each head to reduce the required field of view. The skull and brain were then immersed in fluorinated fluid (Galden, Solvay) to reduce the susceptibility to artifacts and imaged with a 7-T preclinical scanner (Agilent Technologies) employing a 39 mm diameter birdcage radiofrequency coil (Rapid GmbH). 3D FastSpin–Echo (FSE) images were acquired with the following parameters: TE/TR =60/2,000 ms, echo-train-length 8, echo-spacing 15 ms, matrix 192 × 128 × 192, isotropic 150 μm voxel size, and acquisition time 104 min. The images were converted to NIFTI format from the manufacturer's proprietary format using in-house software.

Structural brain images were analyzed by a voxel-based morphometry (VBM) method that allowed automated comparison of the gray matter changes between Low-CR and High-CR rats and Ctr rats. The VBM method was performed as described by Ashburner and Friston (8) with modifications for rodent MRI datasets reported by (7), using statistical parametric mapping (SPM8, http://www.fil.ion.ucl.ac.uk/spm/), FMRIB Software Library (FSL v5.0; http://www.fmrib.ox.ac.uk) and custom-written scripts in MATLAB (MathWorks Inc., Natick, MA, USA), as previously reported (9).

VBM analysis was complemented by a region of interest (ROI) volumetric analysis. We used a set of 115 bilateral ROIs, covering the whole brain, derived by combining the rat brain "Waxholm" (9)and "Tohoku" atlases (10), as described by Kim et al. (2022)(doi.org/10.1101/2022.11.22.517484). For each rat, the standard label image containing all ROIs was back registered to the subject using the inverse deformation field calculated during the VBM analysis. Total brain volume was then calculated by summing the volumes of all 115 subject-registered ROIs.

*In vivo recordings*

Polysomnographic recordings

Electroencephalographic (EEG) and electromyographic (EMG) recordings were performed in rats chronically implanted with electrodes for differential fronto-parietal electrocorticographic (EcoG) derivations and nuchal muscle EMG. Surgeries for EEG implants were performed on head-fixed animals (3 months old) in a stereotaxic apparatus under isoflurane anesthesia (1.5-2.5%, in oxygen) and with eye-protecting gel (Viscotears). Animals received preoperative analgesia via an injection of buprenorphine (0.05 mg/kg, s.c.) and a mix of lidocaine/bupivacaine injected locally under the skin of their heads. EEG electrodes were four gold-plated screws (diameter 1.1 mm) implanted into the skull over the two hemispheres to obtain two fronto-parietal differential derivations; two additional screws were inserted for implant stabilization. In addition, two wires were inserted into the neck muscles for EMG recordings. A headmount female connector (Pinnacle Technology Inc.) was soldered to EEG and EMG electrodes and sealed to the skull with dental cement (Paladur, Heraeus Kulzer). Animals recovered from surgery for one week with paracetamol added to the drinking water (2 mg/ml) in the home cage. Subsequently, rats were tethered to a swivel commutator via a connecting male cable that features a preamplifying element for biopotentials (Pinnacle Technology Inc.). Rats were habituated one additional week to the experimental cage (circular Plexiglass cages, 35 cm diameter) once electrode implants were connected to the tethering cables. Animals that underwent FC were disconnected at each testing day, only for the time of the test. For analysis of undisturbed sleep, polysomnographic acquisitions were performed in 48-h or 12 h-long sessions. For postextinction sleep analysis, polysomnographic recordings were started immediately following extinction training until the next day (24 h). EEG/EMG signals were digitized at 2 kHz with a data conditioning and acquisition system and recorded and analyzed with Sirenia Software (Pinnacle Technology Inc.). Vigilance states (wakefulness, NREMS, and REMS) were manually scored by visual inspection using 4-s epochs according to well-established criteria by an experienced experimenter (11). Briefly, wakefulness was scored as predominant low-amplitude fast activity associated with EMG bursts of movement-related activity. NREMS was characterized by relatively high amplitude delta-frequency (< 4.0 Hz) EEG periods and reduced EMG activity and REMS was characterized by sustained periods of theta (5.0–8.0 Hz) EEG activity associated with minimal EMG activity (muscle atonia). Power spectra were determined using discrete-Fourier transformation between 0.75 and 90 Hz (0.25 Hz bins) for consecutive 4-s epochs.

Fiber-photometry recordings

Endogenous NE levels were monitored in the hippocampus using the fluorescent reporter (G-protein-coupled receptor-activated-based) $GRAB_{NE}$2h, a novel version of the sensor developed by Feng et al. (12). At p90, rats were anesthetized with isoflurane and infected with AAV9-hSyn-NE2h (WZ Bioscience, Inc.), which was infused in the left hippocampus at a volume of 600 nl (titer $5.15 \times 10^{13}$ vg/ml), targeting the DG with the following stereotactic coordinates (from bregma: AP, -6.2; ML, +2.5, DV, -4.2). Viral delivery was performed with a stainless steel injector (200 µm in diameter) connected to a peristaltic pump operating at an infusion rate of 0.1 µl/min. The needle was left in place 10 min after termination of the injection before being slowly retracted from the brain. Following 3-4 weeks to allow viral expression to occur, a second stereotactic surgery was performed to place a fiberoptic cannula (6-mm long fiber of 400 µm diameter, exposed from a zirconia ferrule with a diameter of 2.5 mm, FC_400/430-0.48_6 mm_MF2.5(G)_FLT, Doric Lenses Inc.) approximately 100 µm above the infected area. The fiberoptic cannula was inserted at a 30° angle in the caudal direction to leave sufficient room for the EEG implants to be placed above the forebrain. The fluorescent signal resulting from LED-driven excitation at 465 nm was monitored while the fiber was lowered to the DG. A sudden increase in the level of emitted fluorescence indicated successful placement of the fiber. The fiberoptic cannula was fixed to the skull with light-curing adhesive (iBond, Kulzer GmbH) and flowable composite (KerrHawe SA). In the same surgery session, the EEG/EMG implants were placed as described above.

Fiber-photometry acquisition sessions were limited to 2-3 h to limit photobleaching. Excitation was provided with a blue LED (CLED 465 nm; Doric Lenses Inc) operating in lock-in mode, with a sinusoidal modulation of 211 Hz. The blue LED was coupled to a fluorescence MiniCube (FMC4_IE(400-410)_E(E460-490)_F(500-550)_S, Doric Lenses) that directed the light to the animal via a low autofluorescence 400-µm-thick fiberoptic patchcord (MFP_400/430/1100-0.57_1.5m_FCM-MF2.5_LAF, Doric Lenses Inc.). Changes in bioluminescence were recorded via a photoreceiver (Newport Visible Femtowatt Photoreceiver Module), acquired at 12 kHz and demodulated through a fiber-photometry console controlled with Doric Neuroscience Studio Software (Doric Lenses Inc.). Acquired signals were downsampled with a factor of 100 and saved as csv files. Offline analyses and alignment to the hypnogram were performed with a routine written for Igor 7 (WaveMetrics Inc.). After applying a low-pass filter at 2 Hz, the whole trace was fitted with a 3-term polynomial fit. The original signal was divided by the obtained fit to correct for bleaching and signal artifacts. The percent change in NE level at NREM-REM transitions was measured for transitions in which at least 5 consecutive NREMS epochs were followed by at least 6 consecutive REMS epochs. The difference between the peak of the $GRAB_{NE}$2h signal before the transition and the mean value of the signal at steady state during REMS was considered.

For verification of viral infection with $GRAB_{NE}$2h and fiber optic placement, rats were deeply anesthetized with pentobarbital (150 mg/kg intraperitoneally, Streuli Pharma) and perfused transcardially (4% PFA, 1× PBS, pH 7.4). The brains were extracted and placed in 4% PFA at 4°C for 24 h, incubated for 48 h in a 30% sucrose PBS solution at 4°C, snap-frozen in isopentane and stored at -80°C. Frozen brains were cut into 30 µm coronal sections with a cryostat (Leica) and stored in cryoprotectant medium. Sections were incubated with goat anti-GFP (Abcam, 1:500), followed by anti-goat (AlexaFluor-568; 1:1000), and by DAPI.

### Supplemental Figures

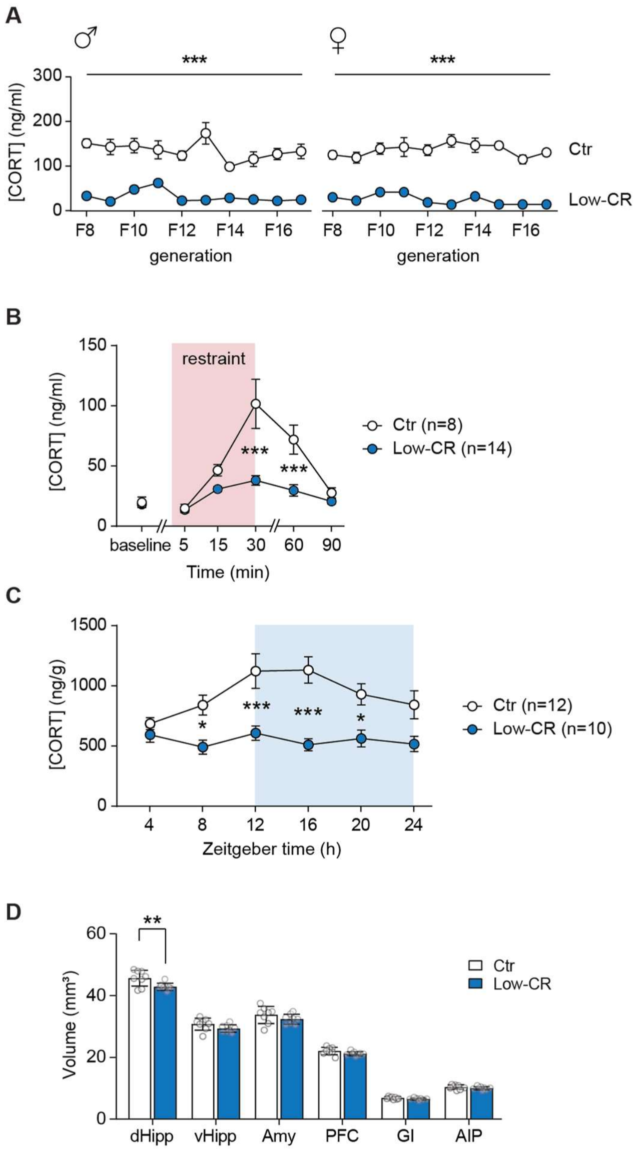


**Figure S1 (related to Figure 1). Plasma CORT levels after repeated stress in rats selected as breeders for Ctr and Low-CR lines and CORT reactivity and morphometric analyses in their progeny.** (**A**) Distribution of plasma corticosterone (CORT) levels in response to stress exposure at the peripubertal stage in outbred Wistar Han male and female rats selected as breeders for the Ctr and Low-CR lines. (**B**) Plasma CORT levels in Ctr and Low-CR rats at baseline and upon exposure to restraint stress (time of exposure indicated by the shaded area). (**C**) CORT levels extracted from fecal boli in Ctr and Low-CR rats across 24 h, with the light phase between 0-12 h of Zeitgeber time. (**D**) Volumetric analysis of brain areas implicated in fear processing. Data are presented as means ± SEMs. Statistical significance was assessed by two-way ANOVA (RM for B and C), with *$P < 0.05$, **$P < 0.01$, and ***$P < 0.001$. DG, dentate gyrus; dHipp, dorsal hippocampus; vHipp, ventral hippocampus; Amy, amygdala; PFC, prefrontal cortex; GI, granular insula; AIP, agranular posterior insula.

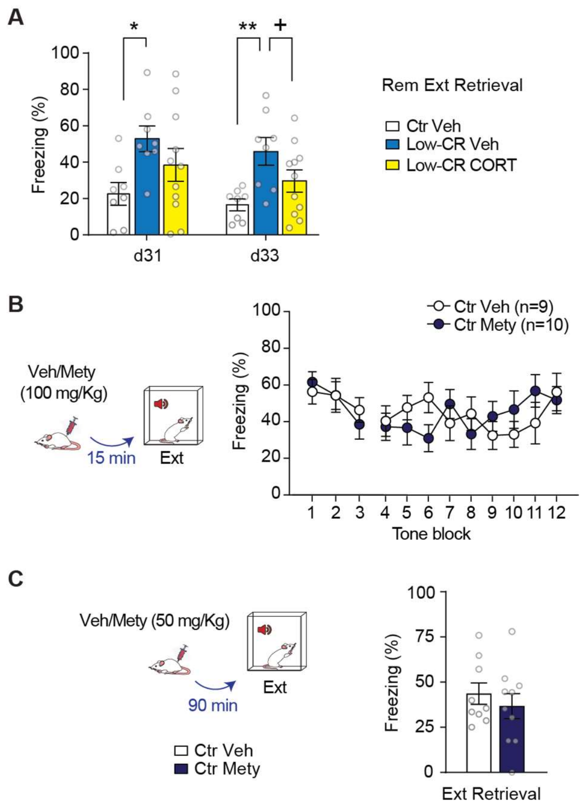


**Figure S2 (related to Figure 3). Complementary data to the HPA axis manipulation experiments.** (**A**) Mean freezing levels during remote extinction retrieval sessions (d31 and d33, experimental timeline in Figure 3) in Ctr and Low-CR rats that had received a vehicle/CORT injection after the extinction training sessions. (**B**) Mean freezing levels during fear recall and extinction of rats injected with vehicle/metyrapone 15 min prior to the extinction training session. (**C**) Mean freezing levels during extinction retrieval in rats that had received a vehicle/metyrapone injection 90 min prior to the extinction training session. All numerical data are presented as means ± SEMs; the number of observations is indicated by datapoints in the bar graphs. Statistical significance was assessed by one-way ANOVA followed by Holm–Sidak's post hoc analysis (A), with $^{+}P < 0.1$, $*P < 0.05$, $**P < 0.01$.

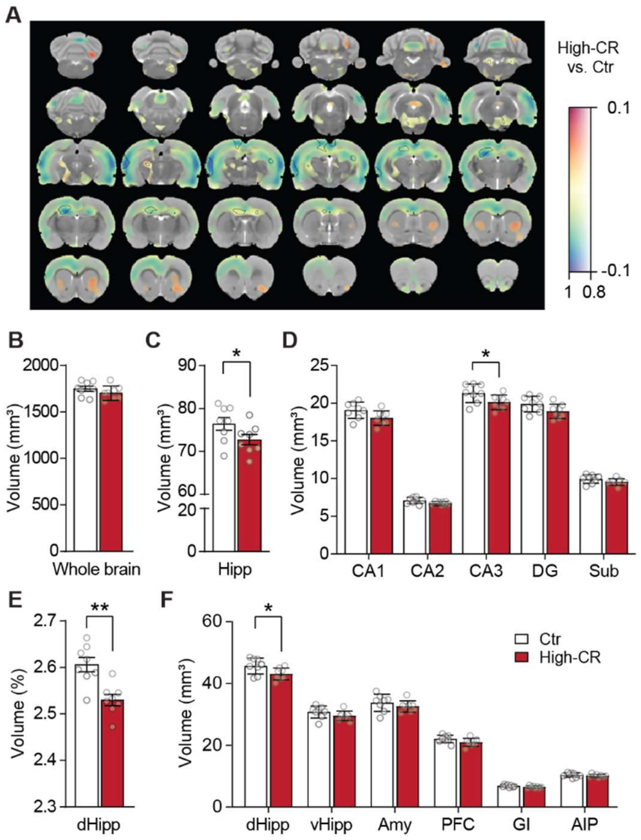


**Figure S3 (related to Figure 4). Higher glucocorticoid responsiveness to stress is associated with hippocampal structural changes.** (**A**) A map of voxelwise differences in GMV between High-CR and Ctr rats calculated from *ex vivo* MR images overlaid on the rat brain MRI template. The colors of the overlay indicate brain volume differences (in $mm^3$) thresholded at 1-p>0.8 (cool colors represent reduced volume, and warm colors increased volume in High-CR compared to Ctr rats), and the transparency of the overlay is related to the *P* value; black contours demarcate regions where uncorrected $*P < 0.01$. (**B** and **C**) Comparison of the volume of the total brain and of the whole hippocampus in Ctr and High-CR rats. (**D**) Volumetric analysis of hippocampal subregions. (**E**) Comparison of dorsal hippocampus volume normalized to the whole brain. (**F**) Volumetric analysis of brain areas implicated in fear processing. All numerical data are presented as means ± SEMs; the number of observations is indicated by datapoints in the bar graphs. Statistical significance was assessed by unpaired *t* test (C-E) or two-way ANOVA (D-F), with $*P < 0.05$, $**P < 0.01$. Nonsignificant comparisons are not indicated. Hipp, hippocampus; CA, cornu ammonis; DG, dentate gyrus; Sub, subiculum; dHipp, dorsal hippocampus; vHipp, ventral hippocampus; Amy, amygdala; PFC, prefrontal cortex; GI, granular insula; AIP, agranular posterior insula.

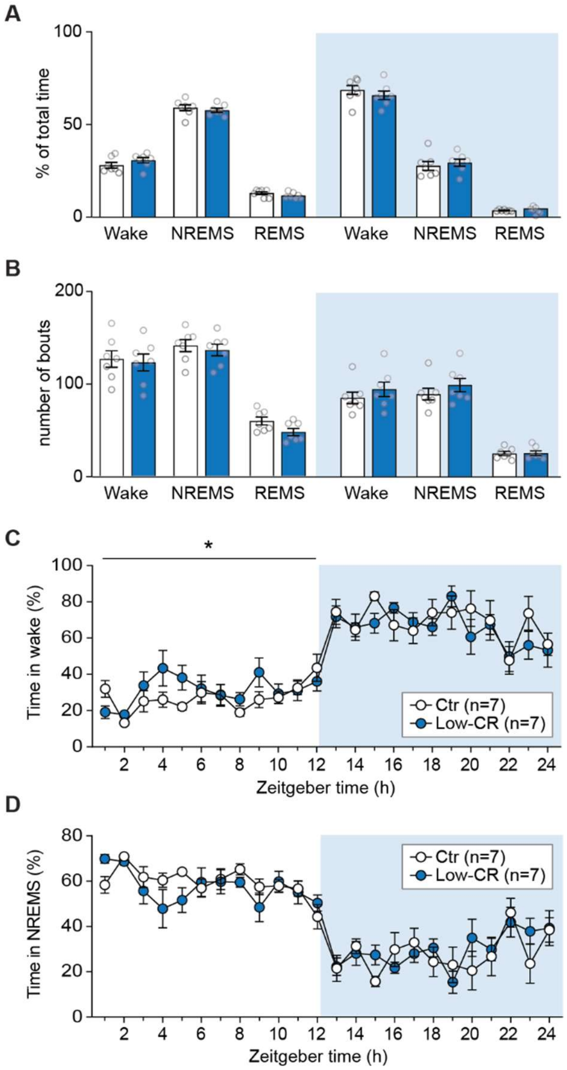


**Figure S4 (related to Figure 5). Blunted glucocorticoid responsiveness to stress is associated with REMS alterations.** (**A**) Percent time spent in different vigilance states averaged during the light and dark phases by Ctr and Low-CR rats. (**B**) Mean number of bouts of wake, NREMS and REMS states during the light and dark phases. (**C-D**) Percent time per hour spent in wake and NREMS states by Ctr and Low-CR rats across the sleep-wake cycle. All numerical data are presented as means ± SEMs; the number of observations is indicated by datapoints in the bar graphs. Statistical significance was assessed by RM two-way ANOVA (C), with *$P < 0.05$. Nonsignificant comparisons are not indicated.

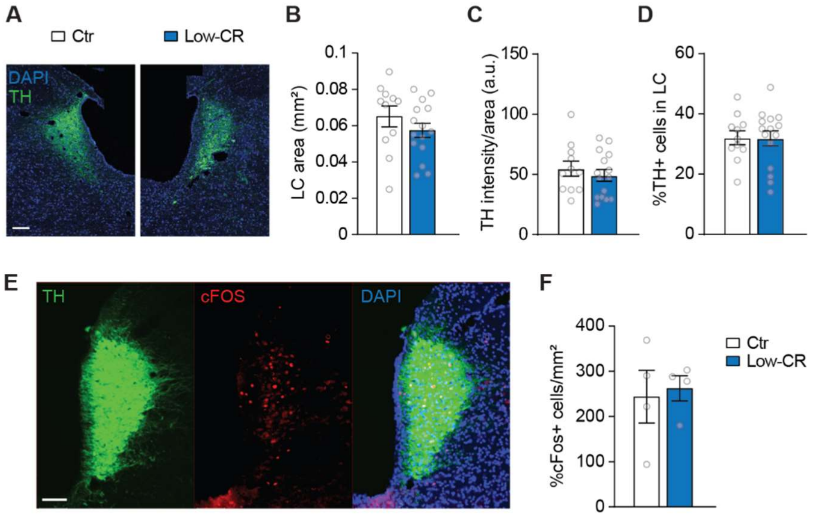


**Figure S5 (related to Figure 5). Lack of differences in markers of LC activity between Ctr and Low-CR rats.** (**A**) Representative images of Tyrosine Hydroxylase (TH) reactivity in LC from a Ctr and a Low-CR rat. (**B-D**) Lack of differences in LC area, TH intensity and percentage of TH+ cells in LC from Ctr and Low-CR rats. (**E**) Representative images of TH and cFos reactivity in a Ctr rats. (**F**) cFos reactivity in LC of rats tested for fear recall after reinstatement. All numerical data are presented as means ± SEMs; the number of observations is indicated by datapoints in the bar graphs. Scale bars = 100 µm.
